# Benchmarking Chemical, Genetic, and Cell Line Encodings for Cancer Perturbation Response Prediction

**DOI:** 10.64898/2025.12.15.694331

**Authors:** Valentyna Zinchenko, Andreas Schlicker, Roman Kurilov, Alison Pouplin, Santiago Villalba, Marc Horlacher

**Author notes:** These authors contributed equally to this work (Valentyna Zinchenko, Marc Horlacher).

## Abstract

Estimating the response of tumor cells to specific perturbations is crucial for identifying effective treatments that selectively target cancer cells while sparing healthy ones, enabling personalized medicine approaches. Large-scale initiatives, such as DepMap, have profiled cancer cell line responses to various drug treatments and gene knockouts, facilitating the development of computational models that predict sensitivity of cancer cells to different perturbations. Existing models utilize diverse methods for encoding perturbations, including various chemical fingerprints and types of gene-gene relationships. They also rely on different architectures and are often trained on distinct datasets. This variability makes it unclear which chemical, genetic, or cell line encoding is most informative for predicting cancer cell viability following perturbation treatment. To address this gap, we systematically evaluated various approaches to encode chemical and genetic perturbations and cell lines on the tasks of predicting cell viability and gene dependency. We found that for genetic perturbations, STRING-based encodings yield the highest performance, considerably outperforming GO-term and protein language model based encodings, which showed promising results in previous perturbation prediction studies. For chemical perturbations, while most encoders showed comparable performance, those pre-trained on other bio-assay data yielded the highest performance. Finally, we found that for cell line encodings, raw gene expression features outperformed more sophisticated approaches, such as transcriptomics foundation model embeddings, as well as genotype-based encodings. Together, our results identify promising approaches for encoding chemical and genetic perturbations and enable virtual screening for perturbations with selective toxicity.

## 1 Introduction

Precision medicine aims to improve patient outcomes by matching the right therapy to the right patient. Precision oncology, in particular, explores unique molecular profiles of tumors, which could harbor selective vulnerabilities, arising from oncogene addiction, tumor-suppressor loss, lineage programs, and metabolic rewiring. Synthetic lethality arising from these vulnerabilities can be used to design customized treatments with higher effectiveness and fewer side effects. Advances in precision oncology drug development are heavily driven by insights from preclinical studies, which mostly use human cancer cell lines. Despite limitations compared to patient tumor samples, cell lines retain many tumor-intrinsic features, while enabling systematic genetic and chemical perturbations at scale, providing a tractable experimental foundation to map vulnerabilities and discover biomarkers that can inform precision oncology (1). Over the past decade, multiple large-scale drug sensitivity screens have been generated, such as GDSC (2) and PRISM (3, 4), profiling viability of multiple cancer cell lines under thousands of compounds, including approved and clinical trial drugs. Meanwhile, gene essentiality of hundreds of cancer cell lines has been mapped using CRISPR/Cas9 or RNAi screens by multiple efforts, such as DepMap consortium (4).

The vast amount of data generated through the afore-mentioned initiatives has been used by multiple efforts to develop machine learning methods that can predict the viability of cancer cell lines, following genetic or chemical perturbations. Such predictive models hold several promises. First, they can enable virtual screening campaigns, allowing researchers to rapidly test novel perturbations against a cell line of interest *in silico*, allowing effective allocation of wet lab resources to the most promising candidates. Second, predictive models can facilitate personalized medicine initiatives by enabling the pairing of patients with treatments through identifying the most appropriate drug for the molecular profile of patients tumor. Finally, interpretation of a model’s predictions via explainability methods can help uncover mechanisms of action, for instance by elucidating why a given cell line is selectively vulnerable towards a given perturbation.

Several studies have leveraged DepMap and PRISM datasets to build machine learning models that capture selective vulnerabilities of cancer cell lines towards genetic and drug perturbations. For instance, Dempster et al. (5) trained per-perturbation Random Forest models and show that gene expression outperforms mutation-derived features for predicting perturbation response. Keyl et al. (6) developed an explainable AI model that accurately predicts treatment responses of chemotherapeutic drugs in unseen tumor samples and identify anticancer potential in repurposed non-cancer drugs. Weiskittel et al. (7) develop a machine learning method based on network biology and demonstrate that, for certain cancer types, gene connectivity can strongly predict cancer vulnerabilities. Related to this, Ben et al. (8) show that biological pathway activity of cancer cell lines is strongly predictive for drug sensitivity. Chiu et al. (9) additionally leverage unsupervised pretraining to build a deep learning model for predicting cancer dependencies, and extend their predictions to 8000 tumors from The Cancer Genome Atlas (TCGA) (10).

To predict perturbation effects, machine learning approaches first have to numerically encode a given perturbation and (unperturbed) cancer cell line. Multiple approaches have been developed to featurize genes, drugs and cell lines, with perturbation prediction models often differing in the encoding strategies they use. Drug encoders range from hand-crafted descriptors and fingerprints (Morgan/ECFP, Mordred) (11–13) to SMILES (14, 15) and graph neural networks (16, 17). Genetic perturbations are commonly encoded through pathway membership (9), GO-term similarity (18), or co-expression (19). To encode cell state, models commonly use gene expression information, either directly or with prior aggregation in terms of pathway activity or gene-set enrichment (5, 8). Finally, foundation models pre-trained on vast amounts of unlabeled data have recently been leveraged to produce modality-specific encodings for a variety of tasks, including perturbation response prediction.

Given the vast number of encoding approaches, together with the heterogeneity of model architectures, it is difficult to discern which gene-gene relationship modality, chemical descriptors or cell state representation is the most informative for predicting cancer cell viability effects for unseen cell lines and perturbations. To address this gap, we systematically evaluate the performance of different encoding strategies on cancer cell viability assay from DepMap (4), both for CRISPR-based genetic knockouts and drug perturbations.

## 2 Results and Discussion

### 2.1 Evaluating Genetic Encoders

In the context of gene perturbation prediction, several gene encoding strategies have been proposed. Roohani et al. (18) propose a GO-term relationship graph, which is subsequently processed by a Graph Neural Network to derive gene-wise embeddings. Klein et al. (20) directly encode gene products by leveraging ESM2 (21) embeddings, a large protein language model, which encodes protein homology and structure. Ahlmann et al. (19) devise a simple linear encoding which, given a *P* × *G* matrix of *P* perturbations and *G* gene expression read-outs, encodes genes *G* by taking the first *N* principle components across *P*. In other words, genes are encoded by their expression changes across a set of perturbations. Towards iteratively designing perturbation experiments via Active Learning, Huang et al. (22) establish gene-gene relationship kernels utilize a number of different modalities, including protein-protein interactions (PPIs), co-occurrence of gene symbols in literature and gene co-expression. Chiu et al. (9) encode genes via sparse binary vector, based on their membership in gene sets derived from the Molecular Signatures Database (23).

#### 2.1.1 Protein interaction and co-expression provide the most informative gene encodings for predicting gene dependency

Based on the existing literature, the following genetic encoders were selected for evaluation: 1) a protein-protein interaction (PPI) network derived from STRING (24), representing physical interactions of gene products; 2) a GO-term (25) gene similarity graph, grouping genes based on similar biological pathways; 3) a biomedical knowledge-graph derived from Ruiz et al. (26), capturing relationships between genes, drugs, diseases and phenotypes; 4) concept vectors obtain from BioConceptVec (27), encoding literature co-occurence of genes; 5) protein language model (pLM) embeddings derived from ESM (21); 6) gene co-expression leveraging GTEx (28) and 7) L1000 (29) datasets, encoding expression changes across tissues and following cell perturbation, respectively.

For each encoding, we trained an instance of a bi-modal neural network on gene dependency data obtained from DepMap (4) (Methods 3.1), schematically outlined in Figure 1A. To ensure that model performance is influenced by genetic encoding alone, cancer cell lines were one-hot encoded, while one the above mentioned gene encoding strategies was used as input to the perturbation encoder. The model was trained to predict gene dependency scores and evaluated cell-line-wise on a set of hold out genetic perturbations via Pearson correlation.

**Fig. 1.**
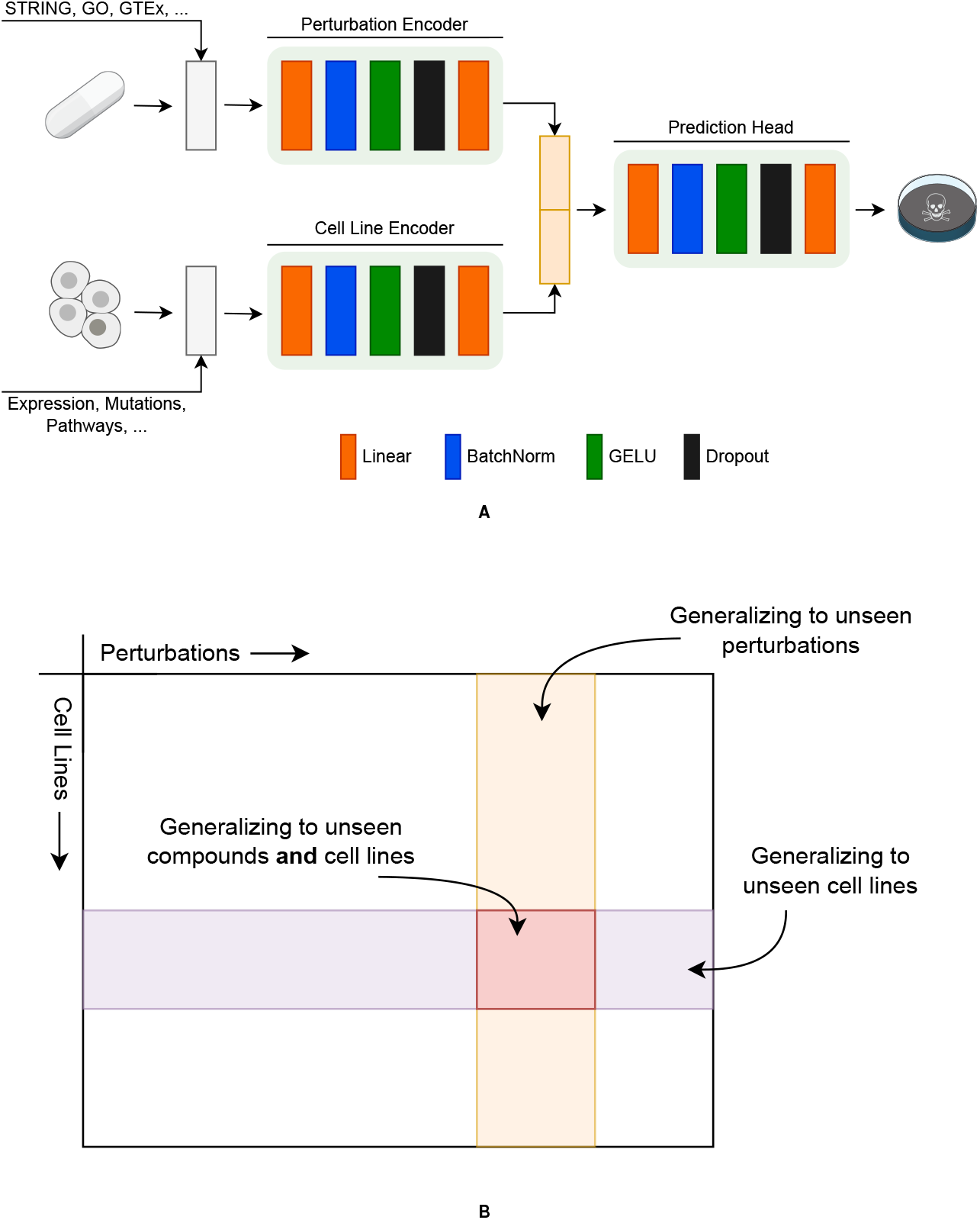
**A** Schematic outline of the bi-modal cancer cell viability predictor. Perturbations (genes or drugs) are first featurized numerically using prior information on gene-gene relationships or chemical descriptors and subsequently embedded by a multi-layer perceptron (MLP) encoder. Similarly, basal (unperturbed) cell states are featurized and embedded with a dedicated cell encoder. The resulting perturbation and cell embeddings are concatenated and fed into a prediction head, which outputs dependency scores for gene perturbations or log-fold-changes for drug perturbations. **B** Data splitting. Generalization performance is evaluated in three regimes — across unobserved perturbations within previously observed cell line, within unobserved cell lines using observed perturbations, and within unobserved cell lines using unobserved perturbation.

All genetic encoders achieved moderate to high Pearson correlation, with a mean Pearson correlation (*mPC*) between 0.57 and 0.74 (Figure 2A), for the GO-term and STRING encoders, respectively. We hypothesize that since STRING not only contains information on a gene’s membership in essential protein complexes and signaling pathways, but also represents *functional* interactions between proteins due to conserved co-expression, co-evolution and text-mining, it provides a strong proxy for gene-gene similarity, which likely translate to similar perturbation effects. The STRING encoder is closely followed by GTEx and L1000 encoders, which achieve mPCs of 0.68 and 0.67, respectively. Similar to STRING, co-expression patterns in GTEx and L1000 matrices likely encode functional relationships via guilt-by-association, thereby representing a powerful modality for predicting gene dependencies. Surprisingly, the knowledge graph (KG) encoder, conceptually operating on a super-set of the PPI network sourced from STRING, yielded markedly worse performance, with a mPC of 0.66. We attribute this to the size of the knowledge graph (108, 447 nodes and 3*M* edges) and featurization via node2vec, which does not discriminate between the heterogeneous relation types in the graph. Together, this may obfuscate relevant substructures in the knowledge graph. Despite their successful use in previous work (18, 20), ESM and GO-term encoders produced a notably lower performance than most alternative encoding methods, with an mPC of 0.61 and 0.60, respectively. In the case of pLM embedding from ESM (21), we hypothesize that while ESM captures protein sequence homology and structural features, these may not directly reflect functional essentiality of a gene (unlike hubs in a PPI graph). Furthermore, genes with similar ESM embeddings may share structural domains but may otherwise participate in unrelated pathways, leading to poor correlation with viability effects. In the case of the GO-term gene-gene similarity graph, unlike relationships in the STRING or knowledge graph, links represent broad functional similarities rather than direct physical or context-specific interactions, which may poorly capture the immediate, mechanistic consequences of gene perturbations. To rule out the case that the low performance of the GO encoder is due to GO graph processing or node2vec featurization, we re-processed the GO-term similarity graph following Roohani et al. (18) (Methods 3.4) and replaced the MLP encoder with a shallow Graph Neural Network. However, this led to a significantly worse performance (Supp. Figure 1), leading us to conclude that GO-term based encoding is indeed inadequate for predicting cell viability following genetic perturbation. Lastly, With an mPC of 0.57, encoding literature co-occurrence of gene symbols showed a lower performance than all aforementioned approaches, indicating that co-mentions in publications are a noisy proxy for gene similarity with respect to essentiality. Despite the marked differences between the encodings, it is important to reiterate that all encoders achieve moderate performance, with a difference between the first and last encoder of 0.17 mPC.

**Fig. 2.**
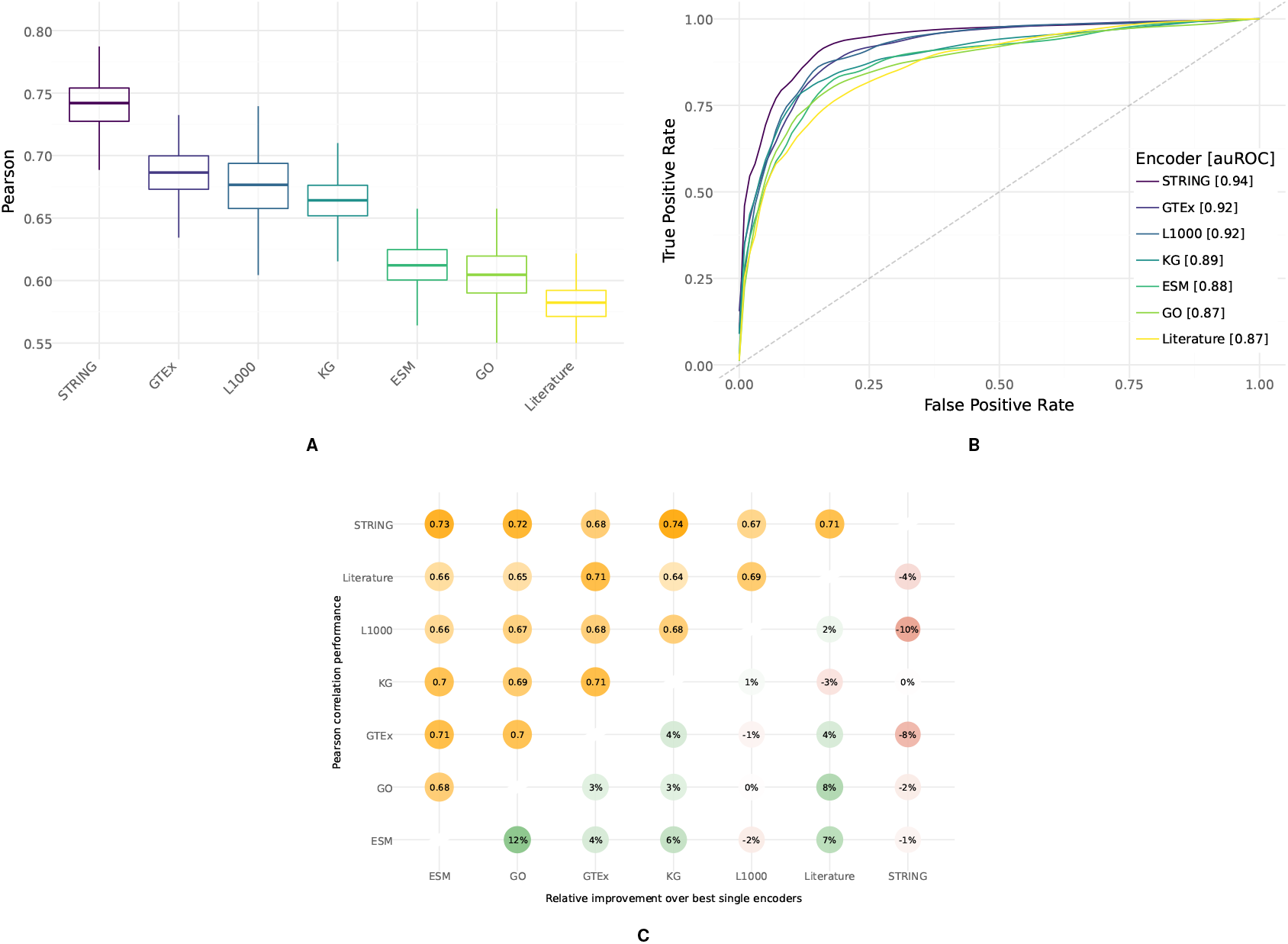
Evaluation of gene encoders. **A** Perturbation generalization performance of gene encoders for each cell line across all hold out genes. **B** auROC performance of gene encoders at a gene dependency threshold of 0.5. **C** Performance of paired gene encoders, measured via mean Pearson correlation (mPC).

#### 2.1.2 Effective Prioritization of Putative Dependencies Across Encoders

In practical applications, virtual cell models are commonly used to prioritize high-confidence targets for experimental validation. To assess the models’ utility in this context, we evaluated their ability to reliably identify putative genetic dependencies. To this end, we binarized DepMap dependency scores into *dependency* / *no dependency* by defining a true dependency as genes with a dependency score > 0.5, following Dempster et al. (30). Subsequently, ROC curves were drawn to assess the models’ ability to rank true dependencies at the top of the predicted dependency score distribution. Figure 2B shows that all encoders can recover putative dependencies with high accuracy, with auROC scores ranging between 0.94 and 0.87 for the STRING and GO/Literature encoders, respectively. This demonstrates the value of the models in terms of prioritizing genetic targets in cancer for experimental validation.

#### 2.1.3 Pairing encoder modalities yields limited performance improvement

Having established that individual genetic encoders can reliably predict cell viability effects of genetic perturbations, we next investigated whether the evaluated encoders contain mutual or potentially synergistic information. Given that genetic encoders represent distinct views on gene-gene relationships, combining encoders may further improve performance. To this end, we exhaustively benchmark pairs of genetic encoders by concatenating both feature vectors before passing them to the encoder neural network. Figure 2C depicts the mPC performance of paired encoder models, together with the relative performance increase over the best single-encoder performance. No pairing improved performance over the single STRING encoder. Strikingly, while this to boost performances for the majority of pairs, pairing STRING with other encoders consistently deteriorates performance. We speculate that since the STRING encoder already has multi-modal properties, encoding gene-gene relationships based on physical PPIs, conserved co-expression and text-mining, further combining it with other encoders provides limited benefit, while the increase in feature dimensionality renders it more susceptible to overfitting and decreases generalization performance.

### 2.2 Evaluating Chemical Encoders

Accurate computational prediction of the effects of an unseen compound on different cell lines may enable virtual drug screening, early removal of toxic compounds, prioritization of compounds for wet-lab testing, and drug repurposing.

Encoding molecular structure and properties in a compact 1D representation is challenging, and multiple approaches have been developed to represent molecules with numeric vectors. Such vectors are used to train computational models that predict molecule properties or biological activities, such as drug effectiveness, toxicity, or ADME properties. Some of the encoding approaches include calculating human-engineered descriptors, such as physicochemical attributes or structural fingerprints that capture connectivity (11, 12), and were used by multiple methods to predict drug sensitivity or post-perturbation gene expression (13, 31, 32). Other commonly used encoding is The Simplified Molecular Input Line Entry System (SMILES) (14), which can be used by drug reponse prediction models either directly (15), or partitioned into substructures like Explainable Substructure Partition Fingerprint (ESPF) (33) and then tokenized (34). Alternatively, SMILES can be used to learn continious data-driven representations in a self-supervised manner, for example, by decoding SMILES or InChI (International Chemical Identifier)(35) from the latent space (36, 37), or using masked language modelling task from the NLP (natural language processing) field (38). Such data-driven representations can be further used for models to predict drug effectiveness or post-treatment gene expression changes (39, 40).

Another way to represent a drug molecule is to use a 2D molecular graph, which could allow to better capture properties relevant to drug interactions. Such graphs can be used directly as input to Message Passing Neural Networks (MPNNs) (16), a subtype of graph neural networks, to encode properties relevant for the final task (17). Recently, multiple efforts have focused on creating MPNN-based foundational models, pre-trained on large numbers of molecular structures and claiming performance gains on various chemical property prediction tasks (41, 42).

#### 2.2.1 Biologically-informed chemical encodings outperform other chemical descriptors

Informed by prior work, we compare the following methods for encoding drug properties: hand-engineered 1) Morgan fingerprints, encoding the topological structure of a molecule (11) and 2) Mordred descriptors that additionally include physiochemical properties (12); 3) CDDD descriptors, learned in a self-supervised way from SMILES representations (37); 4) learned features from MiniMol, a foundational model pre-trained on various quantum-chemical and biological tasks (41); 5) learned features from CheMeleon, a foundational model trained to predict full Mordred descriptors (42); 6) fine-tuned CheMeleon (a trainable MPNN pre-initialized with CheMeleon weights) and 7) a comparable MPNN (16) without prior initialization.

To evaluate these approaches on the task of predicting cell viability we used PRISM drug repurposing resource (3), a pooled drug screening dataset, measuring responses for multiple cell lines upon treatment with various small molecules, which includes compounds either approved, or being investigated for various, mostly oncology-focused indications. We used the same model architecture as for evaluating gene encoders (Figure 1A), categorically encoding cancer cell lines. Cell viability was measured as Log Fold Change, and the evaluation metric was computed cell-line-wise on a set of hold out chemical perturbations.

All benchmarked encoders show moderate performance (Figure 3), with mPCs ranging between 0.25 and 0.47, suggesting that predicting the effects of unseen perturbations is a harder task for small molecules than for gene knockouts, as previously reported in (5). The results show that Minimol (mPC: 0.47), which has partially been pre-trained to predict post-perturbation gene expression data and drug treatment outcomes in high throughput screening bioassays, substantially outperforms all tested encoding methods, highlighting the benefits of using relevant, in-context pre-training tasks. However, in general, deep-learning based featurization, such as fine-tuned CheMeleon (mPC: 0.32) and CDDD descriptors (mPC: 0.25), do not clearly surpass the performance of manually crafted descriptors, such as Morgan fingerprints (mPC: 0.32) and Mordred descriptors (mPC: 0.27). Nevertheless, features derived from a pre-trained CheMeleon model showed 16% higher performance over training a similar MPNN from scratch, while fine-tuning the pre-trained CheMeleon model resulted in an additional 8% improvement.

**Fig. 3.**
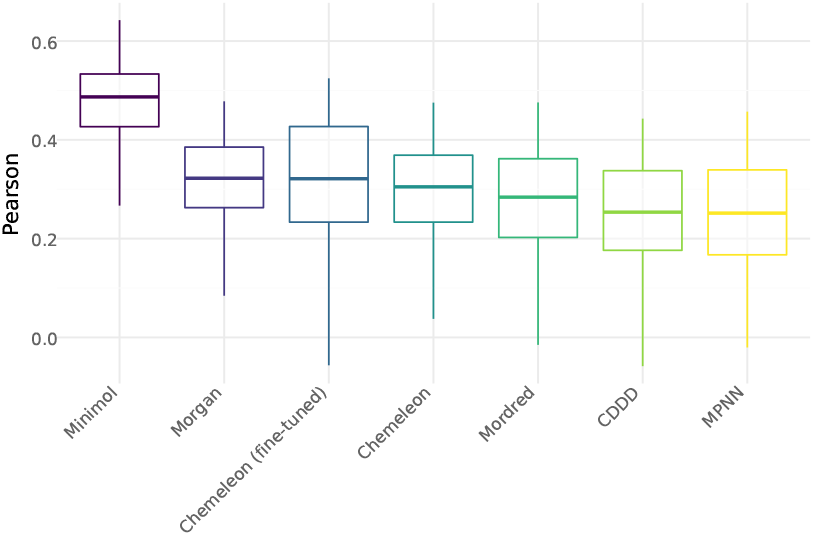
Evaluation of drug encoders. Perturbation generalization performance of chemical encoders for each cell line across all hold out drugs.

### 2.3 Evaluating Cell Line Encoders

Evaluating how well a model can predict the effect of unseen genetic or chemical perturbations allows us to estimate how well the model learned to represent similarities between different perturbations. However, from a precision medicine perspective, a task more challenging is to learn how perturbation outcomes relate to the state of a given cell. Not only would this allow to predict selective vulnerabilities of specific cancer cell lines, but also to better understand the mechanism of these vulnerabilities.

Predicting which cells are sensitive and which are resistant to certain perturbations requires meaningfully encoding the relevant molecular characteristics, representing cell state. A fully comprehensive view of cell state should consider multiple modalities, including gentotype information, gene expression (bulk or single-cell), methylation, proteomics/phosphoproteomics and cell morphology (e.g., Cell Painting). In practice, however, generating all modalities is often infeasible due to tissue constraints, cost, and throughput, so historically both cell lines and patient samples have been profiled primarily by mutation and copy-number data, which have been widely used in predictive models, including for perturbation effects. At the moment, transcriptomics profiling is the de facto standard for representing cell state, and many current pertubation prediction models rely on expression alone (6, 13, 43), or in combination with somatic mutation and copy number variation data (5, 9, 31). The most direct approach uses full single-cell or bulk gene expression profiles as the model input (6, 43), but such high-dimensional, noisy inputs can overfit and generalize poorly to unseen samples, motivating dimensionality reduction and encoding strategies that distill biological signal to improve robustness. Some approaches use a subset of genes; for example, Qi et al. (13) select 5, 000 highly variable genes (HVGs), while Dempster et al. (5) use the top 1, 000 genes whose expression show the highest linear correlation to cell viability scores. Other approaches compress expression information using prior biological knowledge, for example via pathway enrichment analysis (31), or by constructing tumor-specific regulatory networks (44) directly from the data (7). Machine learning-based compression is also common. For example, to reduce gene expression dimensionality, Chiu et al. (9) employ an autoencoder pretrained on TCGA, a large dataset of primary cancer samples (45). A novel promising class of machine learning models, capable of extracting low-dimentional representations, are transcriptomics foundation models pretrained on extensive gene expression datasets. Such models can be used either directly for *in silico* perturbation analysis (46, 47), or as powerful gene-expression-based encoders for cell lines (48).

#### 2.3.1 Raw gene expression outperforms lower-dimensional pathway and latent encodings

To benchmark how well different cell state representations encode cellular characteristics relevant to predicting cell sensitivity to different perturbations, we trained models on chemical and genetic perturbation datasets separately and evaluated how well the effect of observed perturbations can be predicted in unseen cell lines. We compare the following ways of cell line representations: 1) full gene expression and 2) gene expression subset to the most variable genes; 3) hotspot and 4) all damaging mutations; 5) combinations of gene expression and mutation profiles; prior knowledge based encodings, such as gene enrichment on 6) KEGG (40) and 7) Hallmark50 (50 curated, nonredundant gene sets that distill well-defined biological processes and oncogenic states) (23) enrichment, 8) variational autoencoder (SCVI) (50) and foundation models: 9) Geneformer (51, 52) and 10) CancerFoundation (53) (Figure 4A). These representations were used as inputs to the cell line encoder, while the perturbation encoder was replaced by one-hot encoding of the perturbations (either gene knockouts or drugs). Separate models were trained on gene dependency data from DepMap and cell viability from PRISM drug perturbation dataset. The models were evaluated perturbation-wise on a set of hold out cell lines (Figure 4). The majority of perturbations are expected to have the same effect on every cell line, either not causing any change or being universally toxic. Consequently, to better estimate model generalization, only perturbations with selective effect on different cell lines were used for evaluation (described in Section 3.3). However, similar trends are observed when evaluating on the full set of perturbations (Supp. Figure 2).

**Fig. 4.**
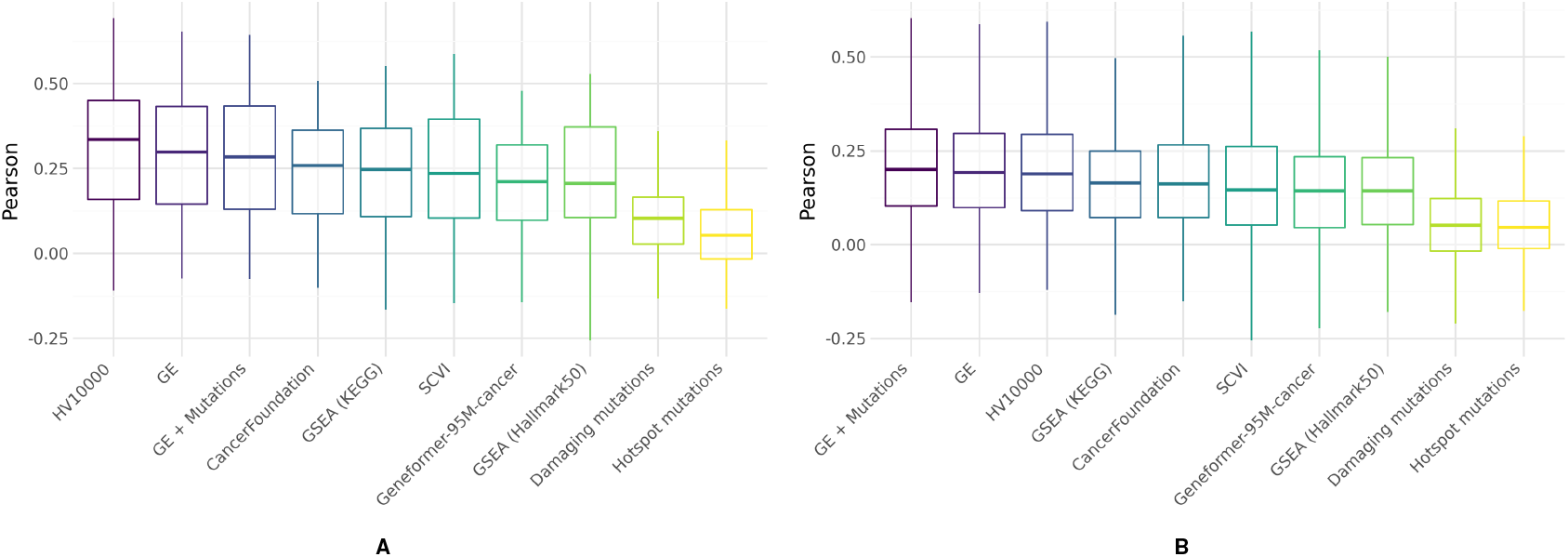
Evaluation of cell line encoders. **A**. Generalization performance of cell line encoders for each highly selective drug across all hold out cell lines. **B**. Generalization performance of cell line encoders for each highly selective gene across all hold out cell lines.

The results confirm that gene expression data predicts perturbation responses more effectively than mutation data. Moreover, combining these two modalities does not bring any notable benefits, suggesting that mutation effects are sufficiently represented in gene expression patterns. Interestingly, for both genetic and chemical perturbations, using gene expression clearly outperforms any prior knowledge or machine learning-based compressed, lower-dimensional representations. Limiting the number of used genes by selecting the most highly variable ones performs on par with the full gene expression, confirming that genes with low variance contain limited information. Expectedly, pathway-enrichment-based features yield better performance when performed on a larger set of pathways, most likely due to failure of the limited set of Hallmark50 pathways to capture nuanced cell states. Strikingly, the tested deep learning-based models, including transcriptomics Foundation Models, did not yield improvements over KEGG-based pathway-enrichment features. We speculate that since foundation models have been trained on single-cell data, they may poorly generalize to bulk RNA-seq data. Additionally, the feature compression induced by average pooling gene-level embeddings may be too restrictive.

## 3 Methods

### 3.1 Data

Gene dependency data from CRISPR knockout (KO) screens were obtained from DepMap (version 25Q2), alongside PRISM drug repurposing data (3, 4, 54). The former dataset contains CRISPR KOs of 17, 916 genes across 1, 183 cell lines, while the later contains perturbation experiments of 6, 790 drugs across 919 cell lines, with a total of 21, 013, 250 and 4, 213, 048 non-empty observations, respectively. For genetic knockouts, the models were tasked to predict gene dependency scores (55), which correct for off-target effects in genetic screens. For drug screens, LFC (Log Fold Change) of perturbed vs. unperturbed cell growth was used. To evaluate cell line encoders on genetic knockouts, during training, genes were filtered to include only those, which are labeled as dependency or no dependency in at least two cell lines. To track performance during training and evaluate the final model’s performance, perturbations and cell lines were randomly split into train, validation and test set at fractions of 0.7, 0.15 and 0.15, respectively. Note that since not all genes are equally covered by all genetic encoders, splitting was first performed on the intersection of genes (covered by all encoders) and the remaining encoder-wise genes were added to the *train* set.

### 3.2 Model Architecture and Training

We use a bi-modal neural network to predict cancer cell viability (Figure 1A). For a given cancer cell line and gene-or drug-perturbation pair, the basal (unperturbed) cell state and the perturbation are first processed separately using a shallow neural network and one of several different cell line and perturbation features (see Sections 3.6, 3.5 and 3.4). Subsequently, the latent cell line and perturbation representations are concatenated and passed to a prediction head to perform the final dependency or LFC score prediction. Unless otherwise specified, the encoders and the prediction head consist of a two-layer multi-layer perceptron (MLP) with batch normalization, GELU activation and dropout. Encoder and prediction head hyperparameter choices for all experiments are listed in Supplementary Table 1. Models were trained using the AdamW optimizer with an initial learning rate of 0.001 and a batch size of 2, 048. Model convergence was monitored on the evaluation set every 200 steps and the learning rate was halved if performance did not improve for 5 evaluations. Models were trained for at most 200, 000 steps, with an early-stopping patience of 15. Evaluation was performed on hold out cell lines when comparing cell encoders, and on hold out perturbations when comparing genetic or chemical encoders.

### 3.3 Model Evaluation

To estimate generalization performance on unseen conditions, Pearson’s correlation coefficient was used. When model performance on unseen cell lines was evaluated, Pearson correlation was computed across perturbations (per cell line), while when evaluating it on unseen perturbations, Pearson correlation was computed across cell lines (per perturbation).

To compare cell line encoders, only perturbations showing highly selective effect on cell lines were used to compute Pearson correlation. For gene knockouts, differential dependencies from (55) were used, defined as selectively essential in some of the cell lines using a six-standard-deviation threshold. For chemical perturbations, highly selective drugs were defined according to (30) as 100 perturbations with the highest NormLRT score (56).

### 3.4 Genetic Perturbation Encoders

#### Co-Expression (GTEx)

Co-expression between genes has long been used for the identification of gene modules via the “guilt-by-association” principle (57). TPM-normalized bulk RNA-seq gene expression data of 19,788 samples, spanning 54 tissues across 946 donors, was obtained from GTEx (28) and processed following Weeks et al. (58). In brief, expression data was log-normalized and scaled gene-wise to a mean of 0 and variance of 1. Then, PCA was performed across genes and the first 256 principle components (explaining >90% of the variance) were selected as the final feature set.

#### L1000 / Connectivity Map

Ahlmann-Eltze et al. (19) proposed to encode gene perturbations based on their co-expression across perturbation conditions in the training dataset and demonstrated that this approach performs on par with state-of-the-art models for post-perturbation gene expression prediction. The L1000, or Connectivity Map, measures the expression of 1, 000 landmark genes across various cell lines and conditions, including gene and drug perturbations (29). L1000 level 5 data was obtained from (29), which yielded differential gene expression signatures across 720, 000 experimental conditions. Similar to GTEx encodings, we compute principle components across conditions and select the first 500, 1, 000 and 2, 000 components as features.

#### Protein Homology (ESM)

Protein language models (pLMs) are ubiquitously used to predict 3D structure and functional properties of proteins, score the impact of missense variants or design new proteins with desired propreties (21, 59–62). Embedding gene products with pLMs offers a direct way to establish gene-gene relationships based on sequence and structural homology of their encoded proteins. Recently, Klein et al. (20) and Adduri et al. (48) leveraged ESM2 (21) embeddings to encode gene knockouts in context of single-cell gene perturbation modeling. Following this approach, ESM embeddings were generated using translations of canonical transcripts for MANE-annotated genes (63). ESM-650M was used to strike a reasonable balance between model expressiveness and embedding dimensionality.

#### GO-Term Similarity

Gene Ontology (GO) term annotations provide a controlled vocabulary to describe the role of gene products in biological processes (25, 64).

For the task of post-perturbation gene expression prediction, Roohani et al. (18) leveraged GO to establish a gene-gene relationship graph by first computing the Jaccard index between two genes *u, v* based on shared GO-terms and then, for each gene *u*, draw edges to the *k* most similar genes. The limitations of this approach arise from the fact that the Jaccard index does not consider the information content of the individual GO-terms, while a threshold on the top-*k* neighbors distorts the scale-free structure of the GO-similarity graph. Instead, we used the GOSemSim R package (65) to compute semantic similarities between genes. Specifically, for two genes *u, v* and their corresponding sets of GO terms *u*_*GO*_, *v*_*GO*_, semantic similarity between all GO term pairs is computed using the Relevance method (66), yielding similarity scores in the range of [0, 1]. Subsequently, semantic similarity scores were aggregated using the Best-Match Average (BMA) strategy. To remove uninformative edges, promote graph sparsity and improve runtime performance, a threshold of 0.5 was applied to the similarity scores, below which no edge was drawn. Note that this is different from thresholding the number of neighbors directly, as it allows for highly connected subgraphs to be preserved.

The final GO similarity graph was then incorporated into the model using two different strategies —as direct input to a shallow Graph Neural Network (GNN), following Roohani et al. (18), and as input to node2vec (67), to project each node to a dense 128-dimensional embedding. The following parameters were used: dimensions = 128, walk_length = 10, num_walks = 400.

#### Protein-Protein Interactions (STRING)

Protein-protein interaction (PPI) networks capture the physical interaction between gene products, thereby encoding pathway and protein complex membership. Further, PPI networks are known to exhibit scale-free structures, with essential proteins acting as networks hubs. Consequently, PPI networks are a promising modality for encoding gene essentiality and dependency. A human PPI network was obtained from the STRING (24) database (v12.0), using the full network with scored links. Besides experimental evidence on physical interaction, this version of the network leverages orthogonal evidence such as co-expression, co-occurrence across species, and text-mining. We then used *node2vec* (67) to featurize the graph into 128-dimensional vectors per gene, using the following parameters: dimensions = 128, walk_length = 10, num_walks = 400. STRING link scores were used as edge weights.

#### Literature

Genes that co-occur in scientific literature tend to be functionally related. For instance, literature co-occurrence of gene products has been shown to be predictive for protein-protein interactions (27). To capture the semantic similarity between genes, we leveraged CBOW word2vec embeddings (68) from BioConceptVec (27).

#### Multi-modal Knowledge Graph

The above-mentioned methods of encoding genes mostly rely on one modality only, such as gene co-expression, protein sequence or physical interaction between proteins. Recently, multiple efforts have focused on integrating heterogeneous sources of biomedical knowledge in networks called knowledge graphs, which connect various entities like genes, proteins, drugs, diseases, phenotypes, pathways, and other types of evidence. Consequently, gene representations learned from such graphs contain information not only about protein co-ocurence or interaction, but also about being involved in similar functions, diseases or causing similar phenotypes. We use the knowledge graph by Ruiz et al. (26), which contains more than 100,000 nodes of different types and, in addition to genes, includes drugs, diseases, biological phenotypes, molecular functions, biological processes and cellular components. The connections between the nodes were derived from various databases including STRING (24), Gene Ontology (64), Drugbank (69), DisGeNet (70) and BioGRID (71). Following Huang et al. (22), we apply *node2vec* (67) on the full knowledge graph to then use the embeddings of gene nodes as gene features. We use the following *node2vec* parameters: dimensions = 128, walk_length = 20, num_walks = 400.

### 3.5 Chemical Compound Encoders

#### Morgan Fingerprints

Morgan, or extended-connectivity fingerprints (ECFP) (11), are hand-engineered molecular descriptors that convert a 2-dimensional graph into a bit vector (commonly 1024–2048 dimensions). They are generated by iteratively enumerating atom-centered neighborhoods up to a chosen radius and hashing these local environments into identifiers. The Morgan fingerprint used here corresponds to ECFP4 (radius 3), generated as a 1024-dimensional bit vector that accounts for chirality. All features were normalized to account for the differing scales of the descriptors.

#### Mordred Descriptors

Mordred (12) is a Python package that generates a collection of interpretable molecular descriptors. Unlike the Morgan fingerprints, which encode only the topological structure of a molecule, the Mordred descriptors also capture their physio-chemical properties. They can be computed from either the 2D molecular graph alone or by incorporating 3D conformer coordinates. In these cases, resulting feature vectors have 1613 and 1826 dimensions, respectively. In this project, both the 2D and 2D+3D normalized representations were evaluated, but only the latter is reported in this paper, as it performed better.

#### Continuous Data-Driven Descriptors (CDDDs)

Continuous and data-driven molecular descriptors (CDDDs) (37)are obtained from an autoencoder trained to translate between different molecular string formats (SMILES to InChI), thereby forcing the model to capture the underlying structure of the molecule. To enrich this representation, additional prediction tasks are used to tune the latent space and incorporate physicochemical information. The model is pretrained on millions of molecules, and once trained, it can encode any SMILES string into a fixed 512-dimensional vector. The CDDD embeddings are not graph-invariant. To ensure consistency, we used RDKit (72) to canonicalize the input SMILES before generating the descriptors.

#### MiniMol

MiniMol (41) is a message-passing GNN that encodes molecular graphs into a 512-dimensional embedding per molecule. It is pretained in a supervised multi-task regime on different endpoints consisting of quantum-chemical properties, high-throughput bioassays, and transcriptomic responses. During pretraining, the encoder and small task heads are optimized jointly. In this paper, the frozen encoder has been used to export the fingerprints.

#### CheMeleon

CheMeleon (42) is a message-passing GNN that encodes molecular graphs into 2048-dimensional molecule embeddings. It has been pretrained on PubChem molecules to predict the full Mordred descriptor vector. The pretrained encoder can be directly used for feature extraction; however, fine-tuning on the end task is recommened. Both approaches were tested in our work.

#### MPNN

In addition to the previously mentioned descriptors, we also evaluated whether chemical compound representations can be learned from scratch for the given task. To this end, we use the ChemProp package (73) to initialize an MPNN following CheMeleon architecture: a BondMessagePassing layer with the following parameters: d_v = 72, d_e = 14, d_h = 2048, depth = 6. This network is trained from scratch, thus not benefiting from transfer learning.

### 3.6 Cell Line Encoders

#### Gene Expression

Gene expression information is readily available and straight-forward to obtain for cancer cell lines and patient tumor tissue dissections. The batch-corrected log-normalized TPM values for all cell lines were obtained from the DepMap portal (4) (version 25Q2). To encode cell state we either used full gene expression vectors (19, 139 genes), or limited it to the top most variable genes across cancer cell lines.

#### SCVI

To further compress gene expression information, we used SCVI, a variational autoencoder framework (50), which projects gene expression vectors into an informative latent space. To obtain cell line features, we trained SCVI on full gene expression data of all cancer cell line models from DepMap, using 3 layers and 500 latent variables. The latent embedding of each cell line was further used as its cell state features.

#### Geneformer

In order to learn more informative latent spaces for single-cell gene expression, Theodoris et al. (51) developed Gene-former, a transcriptomics foundation model that is trained on 30M single-cell transcriptomes using a masked language modeling objective on the relative ordering of genes by expression. Here, we used Geneformer variant 95M-cancer (52) that was trained on a bigger dataset containing cancer cells, which is more relevant for the task at hand. As Gene-former yields an embedding vector for each gene, embeddings were averaged across genes to obtain a final cell-line embedding.

#### CancerFoundation

Following the need for cancer-specific gene expression encoding, Theus et al. (53) developed a single cell transcriptomics foundation model trained exclusively on malignant cells. Although trained on single-cell expression data only, the model showed good performance in encoding cell line bulk gene expression data for a similar task of drug response prediction on GDSC dataset (2). Non-batch corrected lognormalized TPM values were used as input for the model. Class tokens were used as cell line embeddings.

#### Pathway Enrichment

Up-or down-regulation of cellular pathways can render cancer cell lines vulnerable or resistant to certain chemical or genetic perturbations. For instance, the tendency of cancer cells to rely on glycolysis over oxidative phosphorylation for energy (Warburg effect) (74) may effect their susceptibility towards perturbations targeting metabolic pathways. Thus, encoding cancer cell lines based on up-or down-regulation of cellular pathways may be a promising avenue. To this end, we used the *fgsea* (*version 1*.*34*.*2*) Bioconductor package (75) to measure the enrichment of two sets of pathways across DepMap cancer cell lines —the “hallmark” gene sets (23), encompassing 50 well-defined biological states and processes, and KEGG MEDICUS (49), encompassing 658 pathways. Each cell line is then represented by the corresponding vector of normalized enrichment scores.

#### Mutational Signature

Accumulation of gain-or loss-of-function mutations are characteristic in cancer cells. To encode cancer cell lines according to their mutational status, we leveraged DepMap genotype data and create two multi-hot encodings, where the *i*’th bit indicated whether gene *i* harbors a hotspot or damaging mutation, resulting in 532-and 19, 517-dimensional feature vectors, respectively.

## Conclusions

The availability of large-scale perturbation datasets has led to an increase in development of models that aim to predict perturbation outcomes *in silico*. These models utilize different strategies to encode cell line identity, genetic and drug perturbations, making it unclear which representations are most effective for predicting cancer cell viability. To address this, we systematically evaluated different encoding strategies for each modality. We find that STRING-based features substantially outperform alternative genetic encodings, including GO-term similarity and protein language model embeddings, which have been previously demonstrated to perform well on related tasks. Furthermore, we find that while the pairwise combination genetic encoders yields to minor to moderate improvements over the best single encoder for most pairs, it does not improve performance beyond the single STRING-based encoder. While we primarily attribute this to inherent multi-modal nature of STRING-derived gene-gene relationships and the presumably high level of mutual information between different encodings, we believe that dedicated deep learning architectures are required to fully leverage synergies between different genetic encoders. For drug representation, beyond the general advantages of MPNN-based foundation models, we highlight the importance of pre-training on relevant tasks, as evidenced by notably higher performance of MiniMol. Strikingly, for cellline encoding, the use of gene expression as input was not surpassed by any dimensionality-reduction approaches, including pathway-enrichment-derived features and foundation model embeddings. Together, this study represents an important step towards identifying the optimal genetic, drug and cell line encoding strategy for predicting cancer cell viability in response to perturbations. Future work should focus on expanding this evaluation to post-perturbation single-cell gene expression prediction.

## ACKNOWLEDGEMENTS

We thank the members of Machine Learning Research Group at Bayer AG, especially Paula M. Zapata and Vladislav Kim, for the insightful discussions. Further, we would also like to thank Gal Barel for providing her valuable expertise on DepMap and related resources.

## Supplementary Figures

**Supp. Fig. 1.**
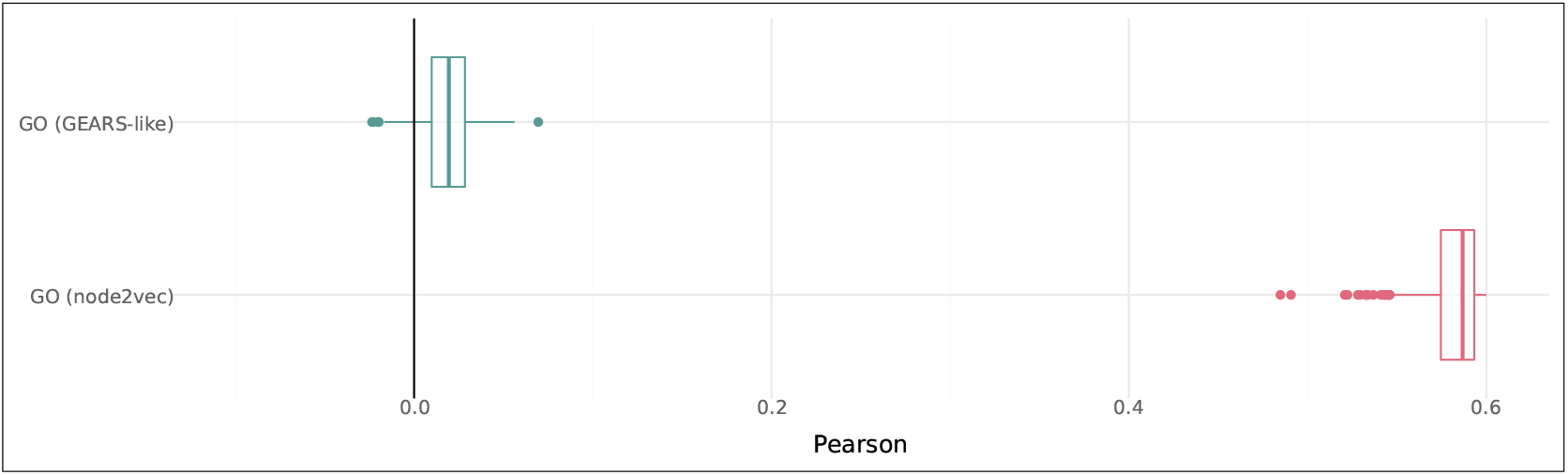
Pearson correlation performance of GO-term encoders using featurization via *node2vec* and a GEARS-like GO-term similarity matrix and GNN encoder (see Methods 3.4).

**Supp. Fig. 2.**
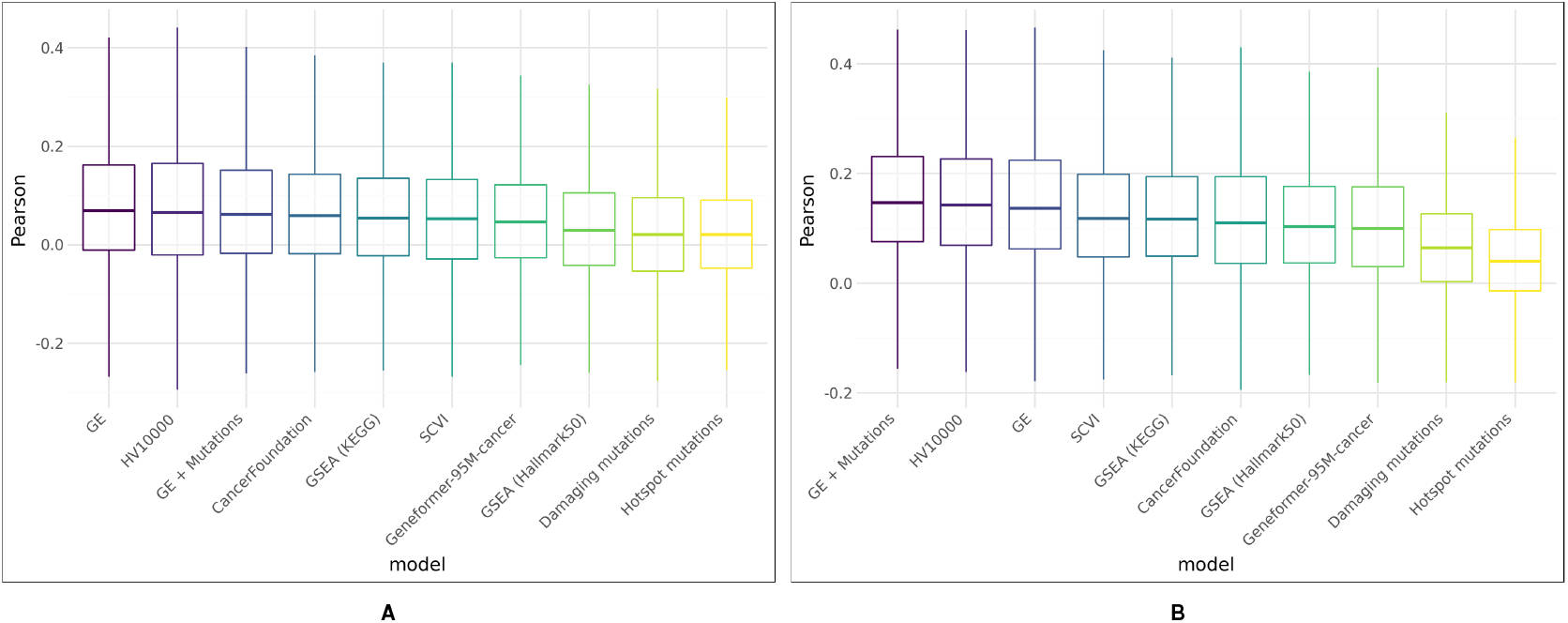
Evaluation of cell line encoders. **A** Generalization performance of cell line encoders for each drug across all hold out cell lines. **B** Generalization performance of cell line encoders for each gene across all hold out cell lines.

## Supplementary Tables

**Supp. Table 1.** Hyperparameter choices for different experiments.

| Evaluation Target | Network module |  |  |
| --- | --- | --- | --- |
|  | Perturbation Encoder | Cell Line Encoder | Prediction Head |
| Genetic | (256, 128) | One-hot | (128, 1) |
| Genetic (Paired) | (256, 256) | One-hot | (128, 1) |
| Chemical | (256, 128) | One-hot | (128, 1) |
| Cell Line | One-hot | (384, 256, 128) | (128, 1) |

